# Antibiotic treatment shapes antigen presentation during chronic TB infection, offering novel targets for therapeutic vaccination

**DOI:** 10.1101/638742

**Authors:** Yu-Min Chuang, Noton K. Dutta, Michael L. Pinn, Chien-Fu Hung, Petros C. Karakousis

**Affiliations:** Department of Pathology, Johns Hopkins University School of Medicine, Baltimore, Maryland, USA; Department of Medicine, Johns Hopkins University School of Medicine, Baltimore Maryland, USA; Department of International Health, Johns Hopkins Bloomberg School of Public Health, Baltimore, Maryland, USA

**Author notes:** Section of Infectious Diseases, Department of Internal Medicine. Yale University School of Medicine, The Anlyan Center for Medical Research and Education, 300 Cedar Street, New Haven, CT 06520-8031. Corresponding author: Dr. Yu-Min Chuang, Department of Pathology, The Johns Hopkins University School of Medicine, 1550 Orleans Street, Baltimore, MD 21231 USA.

**Keywords:** *Mycobacterium tuberculosis*, tuberculosis vaccines, persistence, stringent response, immunotherapy

## Abstract

The lengthy and complicated current regimen required to treat drug-susceptible tuberculosis (TB) reflects the ability of *Mycobacterium tuberculosis* (Mtb) to persist in host tissues. The stringent response pathway, governed by the dual (p)ppGpp synthetase/hydrolase, Rel_Mtb_, is a major mechanism underlying Mtb persistence and antibiotic tolerance. In the current study, we addressed the hypothesis that Rel_Mtb_ is a “persistence antigen” presented during TB chemotherapy and that enhanced T-cell immunity to Rel_Mtb_ can enhance the tuberculocidal activity of the first-line anti-TB drug, isoniazid, which has reduced efficacy against Mtb persisters, C57BL/6 mice and Hartley guinea pigs were aerosol-infected with *Mycobacterium tuberculosis* (Mtb) and, 4 weeks later, received either human-equivalent daily doses of isoniazid alone, or isoniazid in combination with a DNA vaccine targeting *rel*_*Mtb*_. After isoniazid treatment, the total number of Mtb antigen-specific CD4^+^ T cells remained stable in mouse lungs and spleens, as did the number of Rel_Mtb_-specific T cells, although there was a significant reduction in dominant antigen ESAT6-specific CD4^+^ or TB10.4-specific CD8^+^ T cells in the lungs and spleens of mice, Therapeutic vaccination enhanced the activity of isoniazid in Mtb-infected C57BL/6 mice and guinea pigs. When treatment with isoniazid was discontinued, mice immunized with the *rel*_*Mtb*_ DNA vaccine showed a lower mean lung bacterial burden at relapse compared to the control group. Our work shows that antitubercular treatment shapes antigen presentation and antigen-specific T-cell responses, and that therapeutic vaccination targeting the Mtb stringent response may represent a novel approach to enhance immunity against Mtb persisters, with the ultimate goal of shortening curative TB treatment.

## Introduction

Despite the high efficacy of the current 6-month “short-course” combination regimen, tuberculosis (TB) remains a global health emergency. Improper provision and supervision of treatment leads to excess morbidity and mortality, continued transmission, and emergence of drug resistance [1]. The prolonged duration of curative TB treatment is believed to reflect the ability of a subpopulation of *Mycobacteria tuberculosis* (Mtb) bacilli to remain in a non-replicating persistent state in the infected host [2]. These “persisters” exhibit reduced susceptibility to isoniazid [3], which inhibits mycolic acid synthesis in the cell wall [4, 5]. This phenomenon is referred to as “antibiotic tolerance” [6], in which slowly dividing or non-dividing bacteria become less susceptible to killing by antibiotics targeting actively multiplying organisms.

One of the key bacterial pathways implicated in antibiotic tolerance is the stringent response, which is triggered by the rapid accumulation of the key regulatory molecules hyperphosphorylated guanosine ((p)ppGpp) and inorganic polyphosphate (poly(P)) in response to nutrient starvation and other stresses [7]. Mtb has a dual-function enzyme, Rel_Mtb_, which is able to synthesize and hydrolyze (p)ppGpp [8, 9], as well as two polyphosphate kinases (PPK1, PPK2) and two exopolyphosphatases (PPX1, PPX2), which regulate intracellular poly(P) homeostasis [10-13]. The mycobacterial stringent response appears to be a positive feedback loop, as poly(P) phosphorylates and activates the two-component system MprAB, which induces expression of *sigE* and *rel*_*Mtb*_ [14], leading to increased synthesis of (p)ppGpp, which inhibits the hydrolysis of poly(P) by the exopolyphosphatase, PPX2 [13]. Deletion of *Rv2583/rel*_*Mtb*_ results in profound bacterial phenotypes, including defective growth at elevated temperatures, and reduced long-term survival in nutrient starvation and progressive hypoxia [15], as well as in mouse lungs [16]. Rel_Mtb_ deficiency also results in defective Mtb survival in a mouse hypoxic granuloma model of latent TB infection [17] and in guinea pig lungs [18].

Although the Mtb stringent response contributes to the formation of persisters and antibiotic tolerance, it remains to be determined whether this pathway can be targeted therapeutically during chronic Mtb infection. We have shown previously that DNA vaccination with four key stringent response genes (*rel*_*Mtb*_, *sigE, ppk2*, and *ppx1*) generated antigen-specific CD4^+^ T-cell responses and vaccination prior to Mtb aerosol challenge augmented the tuberculocidal activity of isoniazid in mice [19]. However, very limited data are available regarding antigen presentation of stringent response factors during antitubercular treatment *in vivo*, as well as the optimal target for therapeutic DNA vaccination, which is an effective strategy for generating cellular and humoral immunity in various diseases [20]. In the current study, we first characterized the expression of *rel*_*Mtb*_ in Mtb-infected macrophages treated with the first-line antitubercular drug, isoniazid, as well as the abundance of Rel_Mtb_-specific CD4^+^ and CD8^+^ T cells in the lungs and spleens of Mtb-infected C57BL/6 mice following daily oral treatment with human-equivalent doses of isoniazid. Next, we studied the efficacy of DNA vaccination targeting Rel_Mtb_ as an adjunctive therapy to isoniazid in two different animal models of chronic TB infection.

## Material and methods

### Bacteria and growth conditions

Wild-type Mtb H37Rv was grown in Middlebrook 7H9 broth (Difco, Sparks, MD) supplemented with 10% oleic acid-albumin-dextrose-catalase (OADC, Difco), 0.1% glycerol, and 0.05% Tween-80 at 37°C in a roller bottle [21]. An Mtb H37Rv strain deficient in Rv2583c/*rel*_*Mtb*_ (Δ*rel*) and its isogenic wild-type strain [15, 18] were kindly provided by Dr. Valerie Mizrahi and grown as described above in order to validate the Rel_Mtb_ antiserum.

### IC-21 Macrophages infection and real-time PCR analysis

The C57B/6J macrophage cell line, IC-21 (ATCC, No. TIB-186), was grown in RPMI medium with 10% FBS and 1% penicillin/streptomycin. IC-21 cells were divided and plated in a multilayer culture flask (Millipore). At day 0, 10^6^ macrophages were infected with 5×10^6^ of logarithmically growing H37Rv bacilli. The cells were harvested 4 days after infection, RNA was extracted and qPCR analysis was performed, as described previously [19]. The primers were listed in Table 1.

**Table 1.**
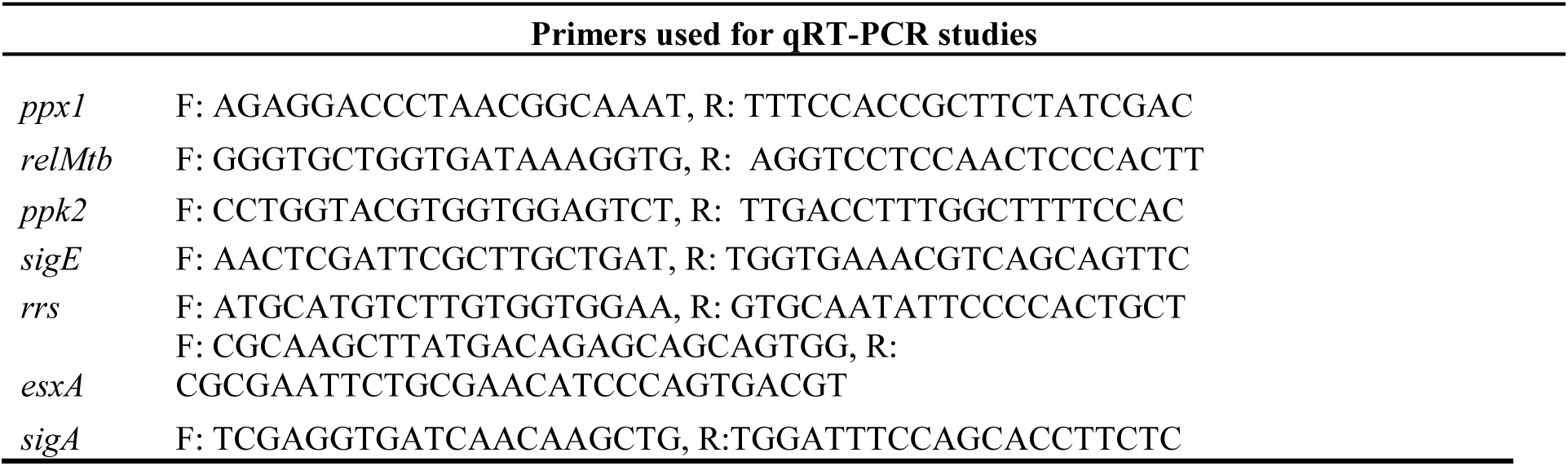
Primers used for RT-qPCR.

### Antigen preparation

The previously generated *rel*_*Mtb*_ expression plasmid, pET15b[*rel*_*Mtb*_] [22], was used for expression and purification of Rel_Mtb_ protein, as previously described [19]. Mtb peptides TB10.4 4 – 11 (IMYNYPAM) and ESAT6 1–15 (MTEQQWNFAGIEAAA)[23] were commercially synthesized (Genescript), dissolved in DMSO and stored at 20°C until use. Mtb whole cell lysate (10μg/ml, BEI) was used to measure Mtb-specific T cells during infection.

### DNA vaccine

The plasmid pSectag2B encoding *rel*_*Mtb*_was used as the *rel*_*Mtb*_ DNA vaccine [19]. ESAT6 was cloned into pSectag2B using the restriction enzymes BamHI and HindIII by forward and reverse primer (F: GCGAAGCTTGTGGCCGAGGACCAGCTCAC, R: CGCGGATCCGCGAACATCCCAGTGACGT). The insertion was confirmed by sequencing. Each DNA vaccine was delivered as previously described [24, 25], and the original plasmid, pSectag2B, was used as control treatment. All procedures were performed according to protocols approved by the Johns Hopkins University Institutional Animal Care and Use Committee. Briefly, each plasmid was delivered by intramuscular injection into the quadriceps femoris muscle of mice (100μg) or guinea pigs (500μg) at the indicated time points.

### Harvest of lung cells and splenocytes

At the indicated time points, the mice were sacrificed and peripheral blood and splenocytes were collected, as previously described [19, 25]. At necropsy, the lungs were perfused with 1ml normal saline by direct injection into the right ventricle of the heart. A random section of the lung was used for cytometry analysis and the tissue samples were incubated in 37° C for 1 hour with intermittent agitation in RPMI medium (Gibco) containing collagenase D (1mg/ml), DNase (0.25mg/ml) and hyaluronidase type V (1mg/ml). The cells were then filtered through a 70-μm nylon filter mesh to remove undigested tissue fragments and washed with complete RPMI medium.

### Intracellular cytokine stain and flow cytometry analysis

To detect antigen-specific T-cell responses by IFN-γ intracellular staining, splenocytes or lung cells were stimulated individually with the purified recombinant proteins, Rel_Mtb_ (10μg/ml), ESAT6 peptide (1 μg/ml) or TB10 peptide (1 μg/ml) for 24 hours at 37°C before the addition of GolgiPlug (BD Pharmingen, San Diego, CA). The concentration of protein and peptides used were according to previous reports [19, 23]. After incubation, the cells were washed once with FACScan buffer and then stained with PE-conjugated monoclonal rat anti-mouse CD4 (BD Pharmingen) and/or APC-conjugated monoclonal rat anti-mouse CD8 (eBioscience). Cells were permeabilized using the Cytofix/Cytoperm kit (BD Pharmingen, San Diego, CA). Intracellular IFN-γ was stained with FITC-conjugated rat anti-mouse IFN-γ and APC-conjugated rat anti-mouse TNFα (BD Pharmingen, San Diego, CA). Flow cytometry was performed by FACSCalibur and analyzed with FlowJo software.

### Aerosol infection of mice with Mtb and therapeutic DNA vaccination

Female C57b/6J mice (6-8 week-old) were aerosol-infected with ∼100 bacilli of wild-type Mtb H37Rv. After 28 days of infection, the mice received isoniazid (10 mg/kg) by esophageal gavage once daily (5 days/week), and were randomized to receive DNA vaccine containing *rel*_*Mtb*_, *ESAT6*, or the empty vector (100 μg) by intramuscular injection. Five mice in each group were sacrificed on Days 30, 60 and 90 after aerosol challenge and the lungs were homogenized and plated for CFU [26]. A partial sample of each infected lung was fixed with 10% buffered formaldehyde, processed, and paraffin-embedded for histological staining with hematoxylin and eosin staining. Morphometric analysis of histology was performed as described previously [27].

### Guinea pig infection and vaccination

Female outbred Hartley guinea pigs (250 to 300 g) were purchased from Charles River Labs (Wilmington, MA). Guinea pigs were aerosol-infected with Mtb H37Rv using a Madison chamber (University of Wisconsin, Madison, WI) to deliver ∼2 log_10_ CFU to the lungs [28]. After 28 days of infection, the guinea pigs received human-equivalent doses of isoniazid (60 mg/kg) daily (5 days/week) by esophageal gavage [29]. During isoniazid treatment, the guinea pigs were randomized to receive either *rel*_*Mtb*_ DNA vaccine (500μg) or empty vector (pSectag2B, 500μg) by intramuscular injection once weekly for four times. 56 days after treatment initiation, three guinea pigs in each group were sacrificed and lung CFU were determined by plating on Middlebrook 7H11 selective agar (BD).

### Enzyme-linked immunosorbent assay and immunoblots

Antigen-specific antibody responses were measured by ELISA as described previously [30], with minor modifications in coating and sera incubation. The 96-well microplate was coated with purified Rel_Mtb_ (1μg/ml) overnight. After blocking, serum from individual vaccinated mice or guinea pigs was diluted 1:100 with PBS, added to the wells and incubated at room temperature for 2 hours. HRP rabbit anti-mouse antibody (Abcam) was used for ELISA with mice sera. For immunoblots, the serum from *rel*_*Mtb*_ DNA-vaccinated mice was used as the primary antibody to detect Rel_Mtb_ expression levels. Monoclonal Anti-Mtb HSP70 (Clone IT-40, BEI) was used to detect the housekeeping protein, HSP70 (DnaK) [19]. To detect anti-Rel_Mtb_ IgG in guinea pigs, Rel_Mtb_ protein was loaded in each lane and proved by probing with anti-His antibody. The serum from vaccinated guinea pigs was used as a primary antibody at 1:50 dilution. HRP-linked goat anti-guinea pig antibody (Novus) was used for ELISA and immunoblots.

### Statistical analysis

Data from at least three biological replicates were used to calculate means and standard error (SEM) for graphing purposes. Pairwise comparisons of group mean values for log_10_ counts and flow cytometry data were made by using one-way analysis of variance and Bonferroni’s multiple comparison test posttest with GraphPad prism 7 (GraphPad, San Diego, Calif.), and a P value of <0.05 was considered significant.

## Results

### Antibiotic treatment shapes antigen presentation and T-cell responses

It has been shown that antitubercular treatment decreases Mtb-specific T cells [31, 32]. Using an intracellular cytokine-releasing assay after stimulation with ESAT6 and TB10.4 peptides, we found that ESAT6-specific CD4^+^ and TB10.4-specific CD8^+^ T cells were significantly reduced in the lungs and spleens of Mtb-infected mice following isoniazid treatment (Figure 1). To further determine if isoniazid treatment can affect antigen-specific T cells, we used Mtb lysate to stimulate splenocytes and lung-derived T cells. Intracellular cytokine staining revealed a significant reduction in Mtb-specific CD8^+^ T cells in the spleens, while Mtb-specific CD4^+^ T cells were increased in the lungs and unchanged in the spleens (Figures 2A and 2B).

**Figure 1.**
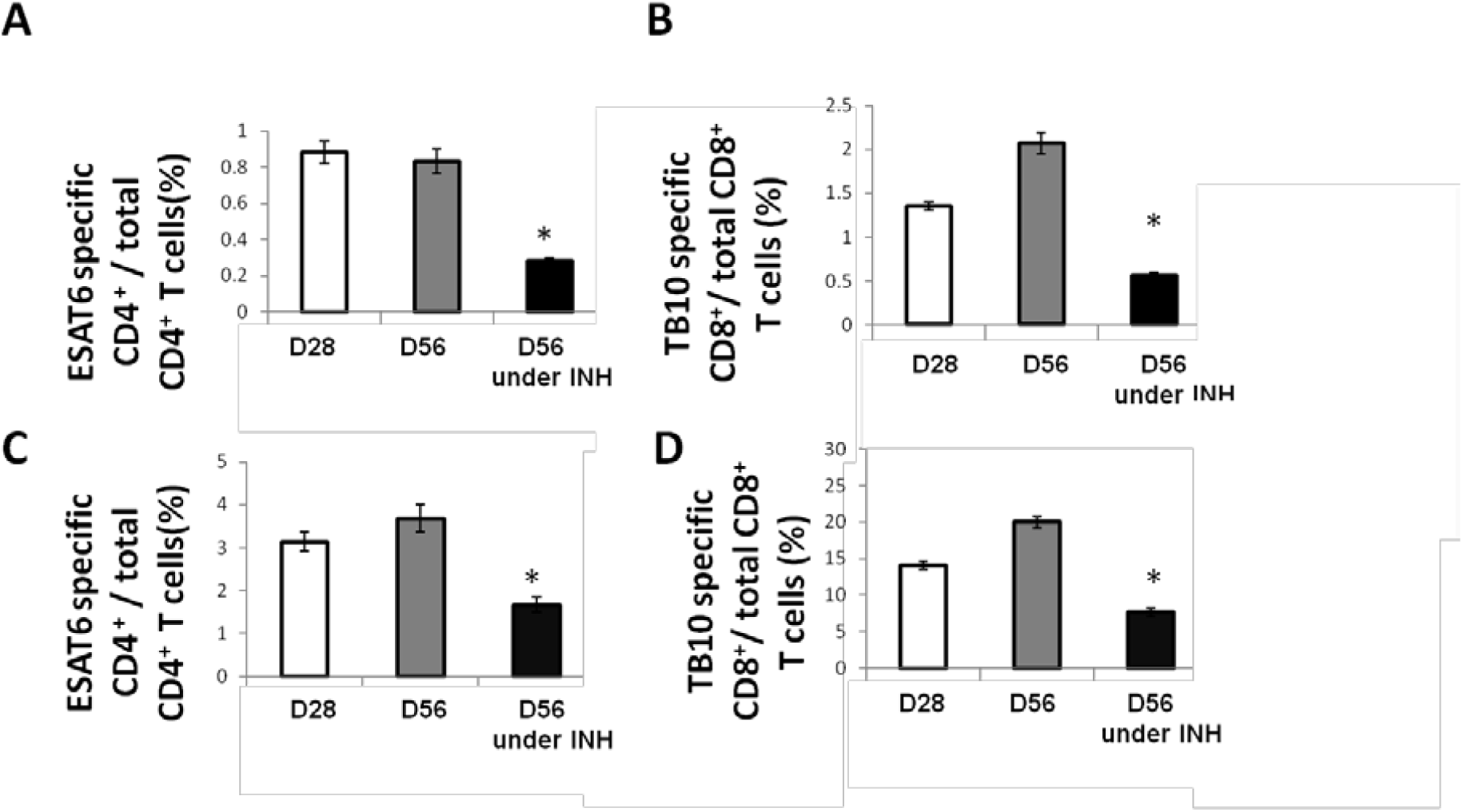
Isoniazid treatment decreases the abundance of immunodominant antigen-specific T cells. C57B/6J mice were aerosol-infected with wild-type Mtb H37Rv and 4 weeks later, one group of mice received daily isoniazid treatment and the control group was untreated. After 4 weeks of treatment, ESAT6-specific CD4^+^ (A) and TB10.4-specific CD8^+^ (B) T cells in the spleen were measured by intracellular cytokine staining after stimulation with ESAT6 and TB10.4 peptides. Lung-derived ESAT6-specific CD4^+^ (C) and TB10.4-specific CD8^+^ (D) T cells were measured by intracellular cytokine staining after stimulation with ESAT6 and TB10.4 peptides. N=3-4, *p<0.05 compared to D56 without isoniazid treatment.

**Figure 2.**
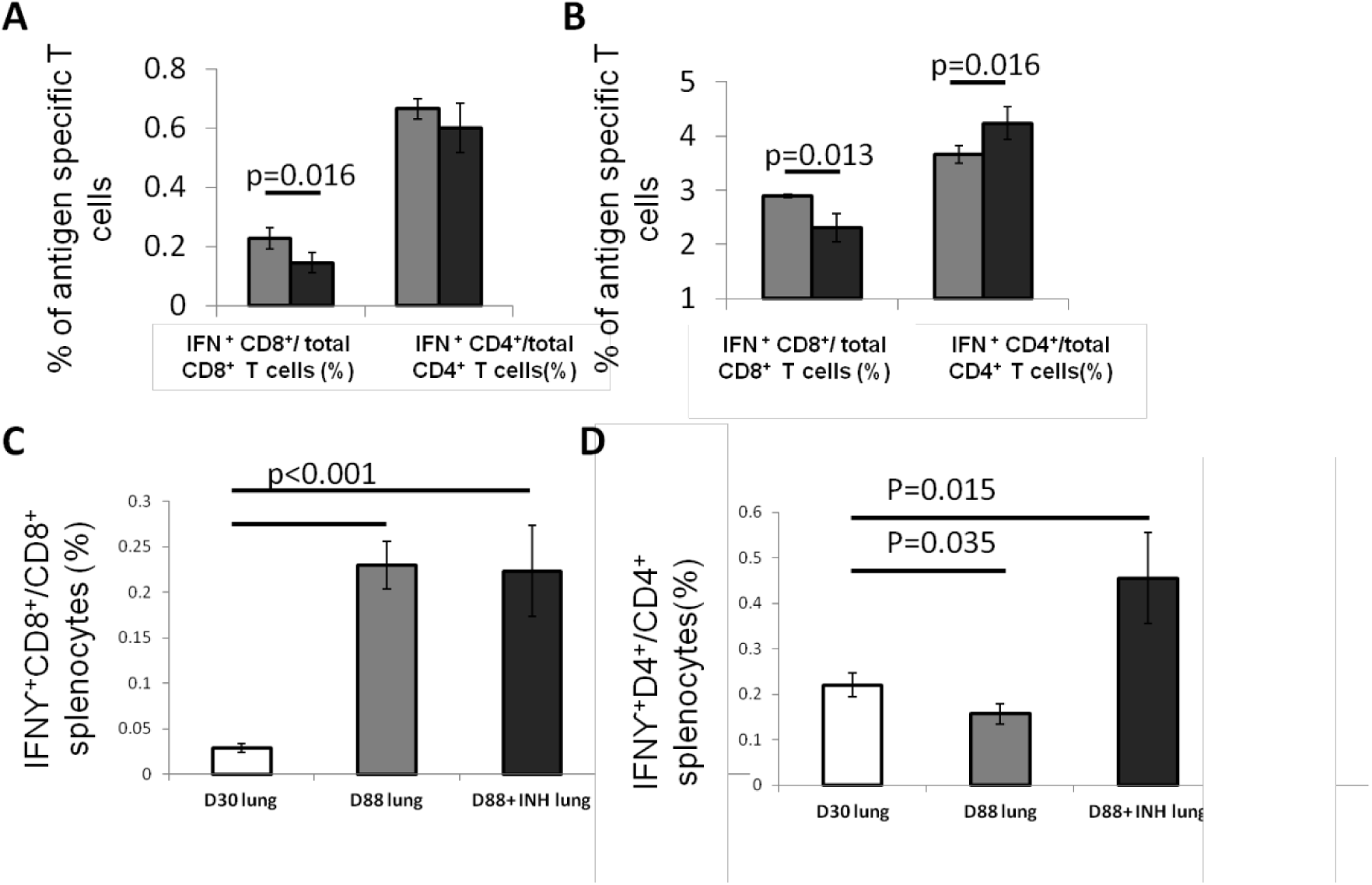
Isoniazid shapes antigen presentation and Mtb antigen-specific CD4+ T cells. C57B/6J mice were aerosol-infected with wild-type Mtb H37Rv and 4 weeks later, one group of mice received daily isoniazid treatment (black bar) and the control group was untreated (gray bar). After 4 weeks of treatment, the splenocytes (A) and lung cells (B) were stimulated with Mtb lysate and antigen-specific CD4^+^ or CD8^+^ T cells were measured intracellular cytokine staining for IFNγ release. N=5. To determine antigen presentation during antibiotic treatment, the lung cells, including APCs, from mice treated with isoniazid at Day 28 and Day 56 and those from untreated mice were co-cultured with splenocytes from Mtb-infected C57B/6J mice harvested 28 days after infection. IFNγ release from CD8^+^ (C) or CD4^+^ (D) T cells was measured by intracellular cytokine staining to determine the antigen presentation. N=3-4.

In order to further characterize the isoniazid-induced changes in Mtb antigen presentation by lung APCs, we harvested lung cells from Mtb-infected C57B/6J mice that were either treated with isoniazid or left untreated (controls), and co-cultured these cells with splenocytes derived from a separate group of C57B/6J mice, which were aerosol-infected with Mtb one month previously. A significantly increased number of IFNγ-releasing CD4^+^ T cells were detected among splenocytes when lung APC were used from the isoniazid-treated group compared to lung APC from the untreated control group (Fig 2C, 2D). Taken together, our data show that isoniazid decreases ESAT6-specific CD4^+^ T cells and TB10-specific CD8^+^ T cells in the spleens and lungs of Mtb-infected mice, but Mtb antigen-specific CD4^+^ T cells were increased in the lungs after treatment, suggesting that isoniazid increases the diversity of antigens presented by APCs.

### Isoniazid induces expression of Mtb stringent response genes during ex vivo and in vivo infection

We found previously that DNA vaccination targeting four genes of the Mtb stringent response showed synergy with isoniazid against Mtb *in vivo* [19]. To determine if stringent response genes were upregulated during isoniazid treatment, we used RT-PCR to study the expression of each of the four genes of our previously reported stringent-response vaccine (*rel*_*Mtb*_, *sigE, ppk2*, or *ppx1*) during Mtb H37Rv infection of IC21 mouse macrophages, which were exposed or unexposed to isoniazid 2 μg/ml for 4 days [33]. As shown in Figure 3A, *rel*_*Mtb*_ was the most highly induced gene following isoniazid exposure (*p*<0.001).

**Figure 3.**
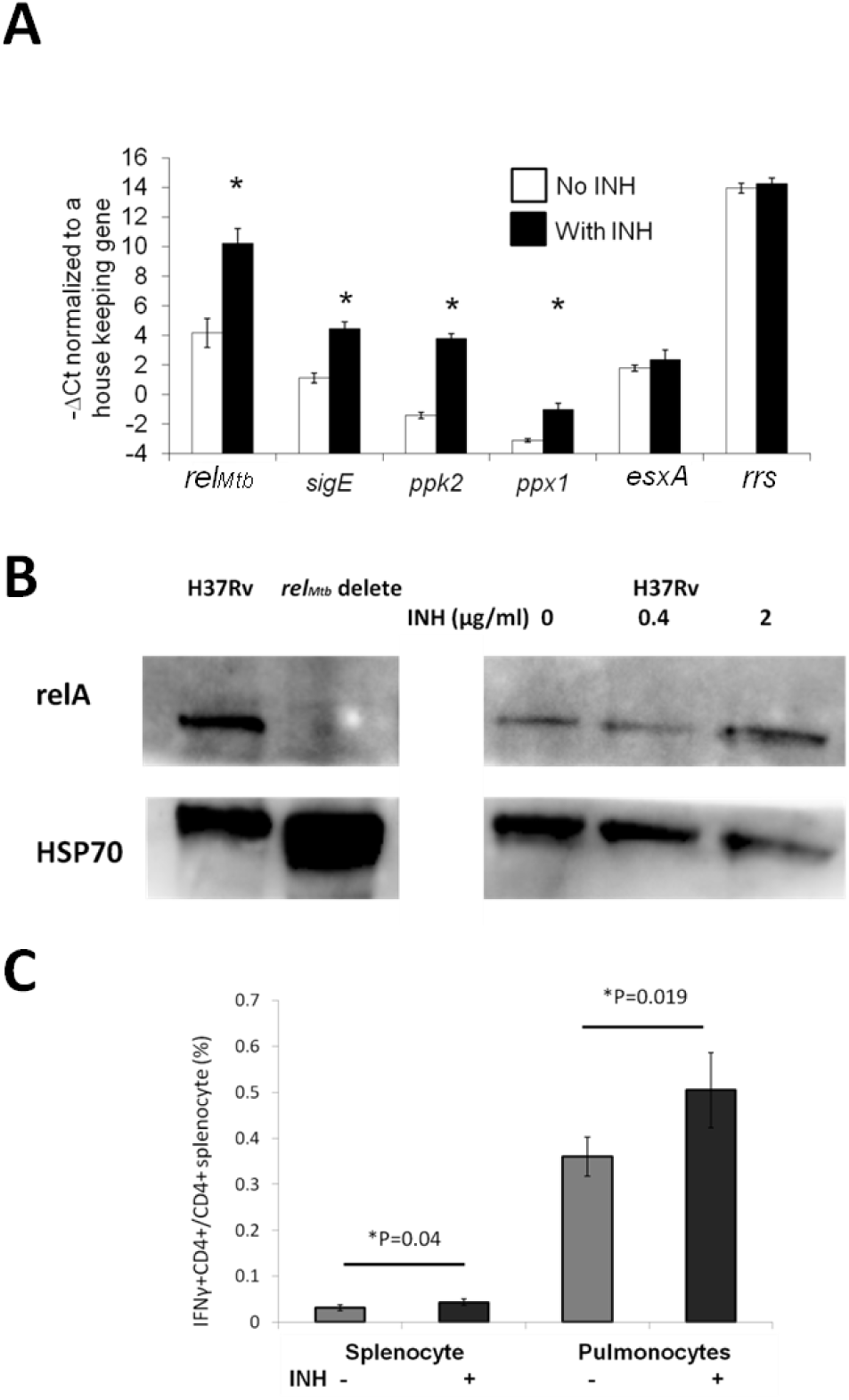
Rel_Mtb_ is upregulated during isoniazid treatment and presented in the lungs of Mtb-infected mice following isoniazid treatment. A. IC-21 macrophages were infected with Mtb H37Rv at an MOI 1:5 and were exposed to isoniazid (2μg/ml) or left unexposed for 4 days. The intracellular bacilli were harvested to analysis expression of the stringent response genes by qPCR. The expression levels of each gene were normalized to that of the housekeeping gene, *sigA*. N=3, *p<0.05 compared to the untreated group. B. Immunoblot confirmation of the specificity of Rel_Mtb_ mouse antiserum and the Rel_Mtb_ expression levels under different concentrations of isoniazid. C. Rel_Mtb_-specific CD4^+^ T cells were significantly increased after isoniazid treatment in the spleens and lungs. N=3-4.

In contrast, expression of the Mtb gene encoding the immunodominant antigen ESAT6 and the housekeeping gene *rrs* did not change significantly after isoniazid exposure of Mtb-infected IC21 mouse macrophages. In order to confirm our findings at the protein level, axenic cultures of Mtb H37Rv were exposed to different concentrations of isoniazid for 4 days. By densitometric analysis, isoniazid exposure (2 μg/ml) increased the expression of Rel_Mtb_ by 2.5-fold relative to unexposed bacteria, following normalization with the housekeeping protein, HSP70 (Figure 3B). Therefore, the stringent response genes were upregulated during infection of macrophages exposed to isoniazid, and Rel_Mtb_ was significantly upregulated following isoniazid exposure during *ex vivo* infection and *in vitro*.

### Rel_Mtb_ antigen is continuously presented during isoniazid treatment in mice

In order to determine the significance of our findings in the context of TB treatment in the mammalian host, we studied Rel_Mtb_-specific CD4^+^ T-cell responses in the lungs of Mtb-infected mice treated with isoniazid as a surrogate for Rel_Mtb_ antigen presentation by APCs during TB treatment. We harvested spleen-derived and lung-derived T cells from Mtb-infected and untreated C57BL/6J mice or infected mice treated with human-equivalent doses of isoniazid for 4 weeks. Consistent with the lower lung bacillary burden in isoniazid-treated mice relative to untreated controls, the proportion of ESAT6-specific CD4^+^ T cells declined in the lungs of the former group relative to those in the latter group. In contrast, the proportion of Rel_Mtb_-specific CD4^+^ T cells was increased in the lungs and spleens of isoniazid-treated mice compared to untreated mice (Figure 3C), suggesting that Rel_Mtb_ antigens continued to be presented during isoniazid treatment.

### Therapeutic DNA vaccination targeting Rel_Mtb_ augments the tuberculocidal activity of isoniazid in C57B/6J mice

Next, we tested whether DNA vaccination targeting the principal stringent response factor, Rel_Mtb_, could augment the bactericidal activity of isoniazid, as in the case of the 4-component stringent-response vaccine [19]. C57BL/6J mice were infected with the virulent Mtb strain H37Rv via aerosol. After four weeks of infection, all mice were treated with isoniazid 10 mg/kg once daily (5 days/week) for four weeks. The mice were vaccinated either with *rel*_*Mtb*_ DNA vaccine (100 μg), or empty vector control (100 μg) once weekly for four weeks starting concurrently with isoniazid treatment. After four weeks of treatment, adjunctive therapy with *rel*_*Mtb*_ DNA vaccine lowered the mean lung bacillary burden by 0.695 log_10_ relative to isoniazid alone (*p*=0.048, Figure 4A). To confirm the therapeutic specificity of the *rel*_*Mtb*_ DNA vaccine, we repeated the mouse study with a longer duration of treatment and added another group of mice, which received isoniazid plus DNA vaccine targeting ESAT6. After 8 weeks of treatment, the group receiving *rel*_*Mtb*_ DNA vaccine again showed a significant reduction in mean lung CFU compared to the empty vector group (*p=*0.045, Figure 4B). In contrast, the mean lung bacillary burden of mice vaccinated with *esat6* DNA vaccine was not statistically different from that of control mice (Figure 4B). Based on these two independent studies, we conclude that immunization with *rel*_*Mtb*_ DNA vaccine potentiates the tuberculocidal activity of isoniazid in mice.

**Figure 4.**
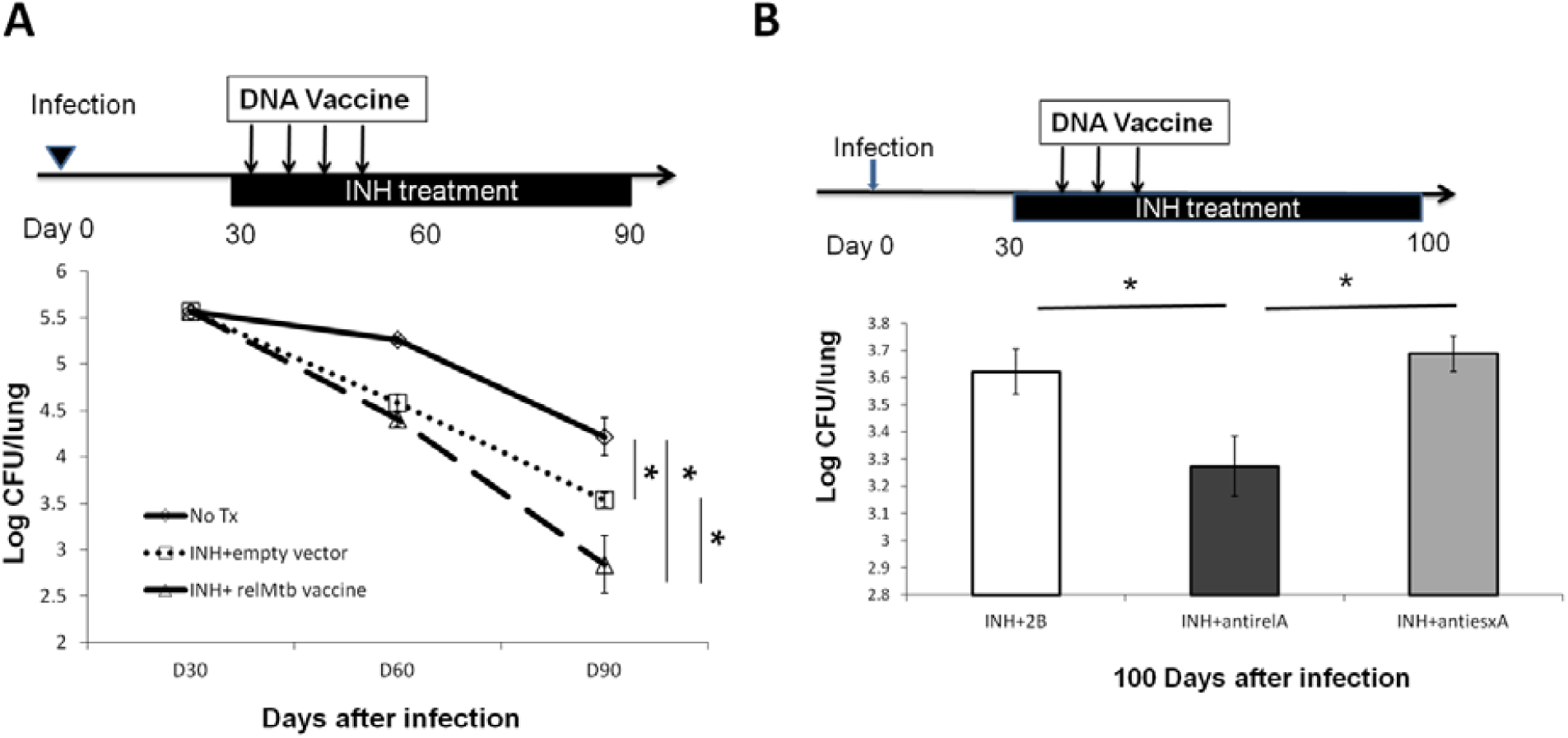
Vaccination targeting Rel_Mtb_ augments the bactericidal activity of isoniazid in Mtb-infected mice. A. C57B/6J mice were aerosol-infected with wild-type Mtb H37Rv and 30 days later, mice were treated with daily isoniazid and immunized with either the *rel*_*Mtb*_ DNA vaccine (100 μg) or the empty vector (100 μg) by intramuscular injection once weekly for four weeks. N=4-5. B. C57B/6J mice were aerosol-infected with wild-type Mtb H37Rv and 30 days later, mice were treated with daily isoniazid for 8 weeks. Concurrent with isoniazid treatment, separate groups of mice received 100 μg of the empty vector (control), *rel*_*Mtb*_ DNA vaccine, or *ESAT6* DNA vaccine by intramuscular injection once weekly for three weeks. N=4-5.

### rel_Mtb_ DNA vaccination reduces Mtb regrowth upon discontinuation of isoniazid treatment

To further study if *rel*_*Mtb*_ DNA vaccine can target persistent bacilli, C57BL/6J mice were aerosol-infected with Mtb and, 4 weeks later, were treated daily with isoniazid for 4 weeks. At the completion of isoniazid treatment, separate groups of mice received either *rel*_*Mtb*_ DNA vaccine (100 μg) or empty vector (100 μg) by intramuscular injection once weekly for two weeks (Figure 5A). Four weeks after cessation of isoniazid, the mean lung bacillary burden of mice receiving the *rel*_*Mtb*_ DNA was lower by 0.5 log_10_ compared to that of the empty vector vaccine group (*p*=0.02, Figure 5B), suggesting that *rel*_*Mtb*_ DNA vaccine can restrict regrowth of Mtb after cessation of antibiotic therapy.

**Figure 5.**
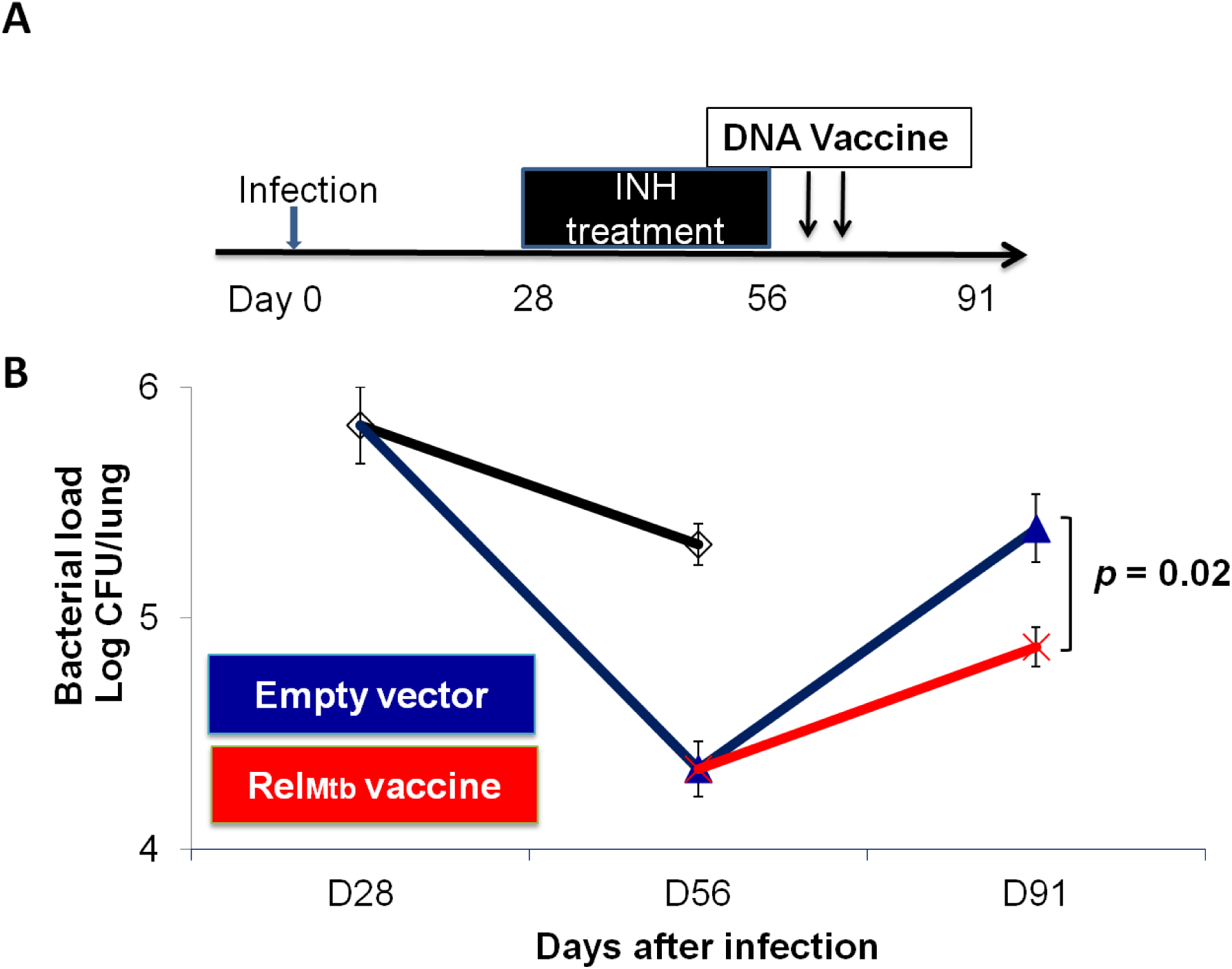
The *rel*_*Mtb*_ vaccine restricts Mtb regrowth following discontinuation of isoniazid treatment. A. Experimental scheme. C57B/6J mice were aerosol-infected with wild-type Mtb H37Rv and 4 weeks later, mice were treated with daily isoniazid for 4 weeks, and treatment was discontinued. Immediately upon discontinuation of treatment, separate groups of mice received either 100μg/ml empty vector DNA vaccine or the *rel*_*Mtb*_ DNA vaccine by intramuscular injection once weekly for two weeks. B. Lung bacillary burden (log_10_ CFU) at day 28, 56 and 91 after infection. N=4-5, *p<0.05.

### rel_Mtb_ DNA vaccine enhances the tuberculocidal activity of isoniazid in guinea pigs

To determine if the adjunctive antitubercular activity of the *rel*_*Mtb*_ DNA vaccine can be duplicated in different species, we infected guinea pigs with virulent Mtb strain H37Rv via aerosol, and 4 weeks later, they were treated once daily with human-equivalent doses of isoniazid (60 mg/kg) for a total of 4 weeks [29]. Separate groups of guinea pigs were vaccinated either with *rel*_*Mtb*_ DNA vaccine (500 μg) or empty vector control (500 μg) once weekly for four weeks beginning with the initiation of isoniazid treatment (Figure 6A). To confirm the immunogenicity of the *rel*_*Mtb*_ DNA vaccine, we used ELISA and immunoblots to demonstrate that this vaccine generated Rel_Mtb_-specific IgG responses (Figures 6B and 6C). At the time of treatment completion, the mean lung bacillary burden in the *rel*_*Mtb*_ DNA vaccine group was lower by ∼0.5 log_10_ relative to the that in the isoniazid alone control group (p=0.05; Figure 6D), demonstrating that, as in mice, *rel*_*Mtb*_ DNA vaccine potentiates the antitubercular activity of isoniazid in guinea pigs.

**Figure 6.**
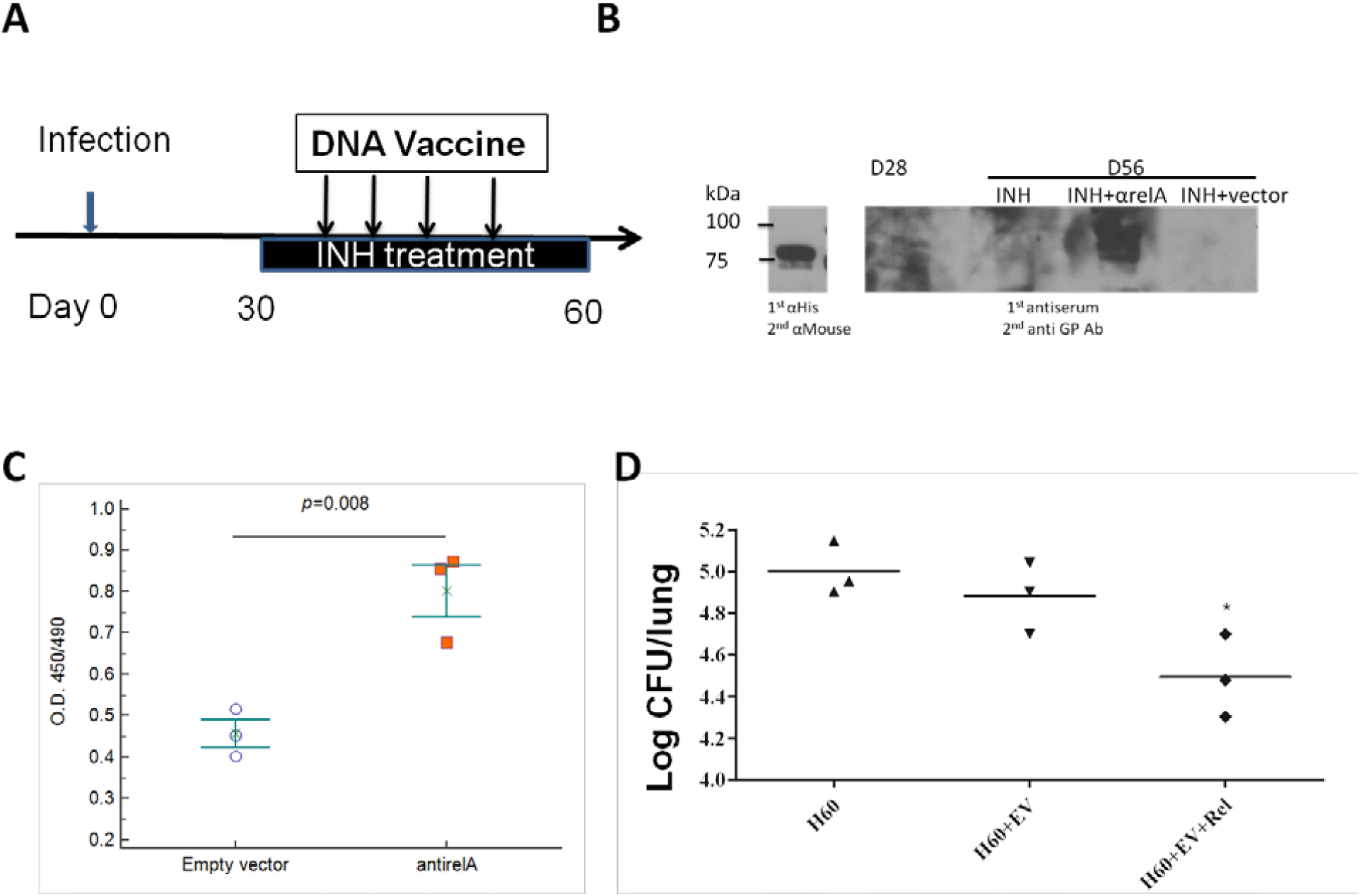
*rel*_*Mtb*_ DNA vaccine enhances the bactericidal activity of isoniazid in Mtb-infected guinea pigs. A. Experimental scheme. Guinea pigs were were aerosol-infected with wild-type Mtb H37Rv and, 4 weeks later, separate groups received daily isoniazid treatment (60 mg/kg) with either empty vector or *rel*_*Mtb*_ DNA vaccine (500μg) by intramuscular injection once weekly for a total of four weeks. B. Immunoblot confirmed the presence of anti-Rel_Mtb_ IgG in guinea pigs vaccinated with *rel*_*Mtb*_ DNA vaccine. C. ELISA detection of Rel_Mtb_-specific IgG responses from groups receiving the empty vector or *rel*_*Mtb*_ DNA vaccine. D. Mean lung bacillary burden (log_10_ CFU) at day 60 after infection (after 4 weeks of treatment). *p<0.05.

## Discussion

Several potential reasons may explain the limited efficacy of immunodominant antigen-specific T cells in eliminating bacteria during chronic Mtb infection, including the reduced amount of immunodominant antigens presented, resulting in a reduction in the number of immunodominant antigen-specific CD4^+^ cells, as well as the continuous release of these antigens, leading to exhaustion of antigen-specific CD4^+^ T cells [34]. Therefore, subdominant Mtb antigens, including those induced during drug treatment, may represent better targets for therapeutic vaccines. In the current study, we show that treatment with the classical bactericidal drug isoniazid increases the abundance of CD4+ T cells specific to the Mtb stringent response factor Rel_Mtb_ in the lungs of mice chronically infected with Mtb, but not those specific to the Mtb immunodominant antigen ESAT6. Furthermore, a DNA vaccine targeting Rel_Mtb_, but not one targeting ESAT6, potentiated the antitubercular activity of isoniazid in two different animal models of chronic TB infection.

Recently, there is significant interest in host-directed therapies to treat TB [35]. Although BCG vaccination does not enhance the bactericidal activity of chemotherapy in the murine model [36], a DNA vaccine expressing heat shock protein 65 has been shown to synergize with conventional antitubercular drugs, further reducing the bacterial burden in the lungs of Mtb-infected mice or non-human primates [36, 37]. A fragment whole cell lysate therapeutic vaccine, RUTI, has shown efficacy in generating protective immunity in pre-clinical studies [38, 39]. However, the primary factors responsible for TB immunity during antibiotic treatment remain unknown. Previously, we have shown that a DNA vaccine targeting key Mtb stringent response factors generates IgG and antigen-specific CD4^+^ T cells, and increased the antitubercular activity of isoniazid when given as a therapeutic vaccine in Mtb-infected mice [19]. The findings of the current study corroborate our previous findings and highlight the importance of immunity to Mtb stringent response factors during isoniazid treatment. One potential explanation for this phenomenon is that the population of persistent bacilli in untreated, chronically infected mice is relatively small [40], but exposure to isoniazid further drives the formation of drug-tolerant persisters [29, 41]. In favor of this hypothesis, isoniazid exposure induces Mtb expression of Rel_Mtb_, which is required for persister formation [7, 42, 43]. Further studies are required to determine the potential utility of the *rel*_*Mtb*_ vaccine as an adjunctive therapeutic intervention in shortening the duration of treatment for active TB or latent TB infection.

Based on our co-culture results, we conclude that isoniazid treatment increased APC presentation of Rel_Mtb_ in the lungs of mice (Figures 2C and 2D). One potential explanation for this finding is that isoniazid increased the amount of Rel_Mtb_ antigen presented due to bacillary lysis. Alternatively, isoniazid treatment may have directly modified Mtb protein expression, leading to altered antigen processing and presentation. We found that ESAT6-specific CD4^+^ and CD8^+^ T cells were significantly decreased in the lungs, even though Rel_Mtb_-specific CD4^+^ T cells were increased after isoniazid treatment (Figure 1), suggesting that isoniazid treatment shapes the diversity of antigen presentation by APCs, enhancing the processing and presentation of subdominant antigens such as Rel_Mtb_.

The *rel*_*Mtb*_ DNA vaccine potentiated the killing activity of isoniazid against chronic TB infection in C57B/6J mice, which develop cellular lung granulomas lacking central necrosis, as well as in guinea pigs, which develop a potent DTH response and more human-like granulomas with necrosis and tissue hypoxia. Interestingly, the adjunctive antitubercular activity of the *rel*_*Mtb*_ DNA vaccine was more pronounced in the guinea pig model, in which the defective survival phenotype of a *rel*_*Mtb*_-deficient Mtb mutant was significantly accelerated relative to the C57B/6J mouse model [44, 45]. These findings suggest that tissue necrosis is not a prerequisite for vaccine efficacy, but its presence may further enhance the activity of isoniazid. Further studies are needed to test the *rel*_*Mtb*_ DNA vaccine in other animal models with necrotic granulomas, including the C3Heb/FeJ mouse [27] and the nonhuman primate [46].

In conclusion, we have shown that isoniazid treatment shapes the diversity of antigen presentation during chronic TB infection in mice, leading to altered antigen-specific CD4^+^ T-cell profiles in the lungs. In addition, this is the first study to use the Mtb stringent response protein Rel_mtb_ as a therapeutic vaccine target. Additional studies are needed to test the therapeutic efficacy of this vaccine in other animal models, and to characterize the immunological basis for its efficacy in mice and guinea pigs.

## Author contributions

Y-M.C. contributed to the design of the study, performance of the experiments, data interpretation and the writing of the manuscript; N.K.D. contributed to the performance of the experiments; M.L.P. contributed to the performance of the experiments; C-F.H. contributed to the design of the study and data interpretation; and P.C.K. contributed to the design of the study, data interpretation and the writing of the manuscript.

